# Multiple RNA regulatory pathways coordinate the activity and expression pattern of a conserved germline RNA-binding protein

**DOI:** 10.1101/2021.01.15.426877

**Authors:** Mennatallah M.Y. Albarqi, Sean P. Ryder

## Abstract

RNA regulation is essential to successful reproduction. Messenger RNAs delivered from parent to progeny govern early embryonic development. RNA-binding proteins (RBPs) are the key effectors of this process, controlling the translation and stability of parental transcripts to control cell fate specification events prior to zygotic gene activation. The KH-domain RBP MEX-3 is conserved from nematode to human. It was first discovered in *Caenorhabditis elegans*, where it is essential for anterior cell fate and embryo viability. Here, we show that *mex-3* mRNA is itself regulated by several RBPs to define its unique germline spatiotemporal expression pattern. We also show that both poly(A) tail length control and translational regulation contribute to this expression pattern. Though the 3’UTR is sufficient to establish the germline expression pattern, we show that it is not essential for reproduction. An allelic series of 3’UTR deletion variants identifies repressing regions of the UTR and show that the expression pattern is not precisely coupled to reproductive health. Together, our results define the pathways that govern the spatiotemporal regulation of this highly conserved germline RBP and suggest that redundant mechanisms control MEX-3 function when RNA regulation is compromised.

## Introduction

Regulation of mRNA metabolism occurs in all cells in all kingdoms of life. In the nucleus, pre-mRNA undergoes splicing, 5’-capping, and 3’-end cleavage and polyadenylation (Shatkin & Manley, 2000). Once mature, mRNA is exported to the cytoplasm where it undergoes further post-transcriptional modifications prior to translation. In the cytoplasm, mRNA can be stabilized by additional poly-adenylation or targeted for degradation by exonucleases through deadenylation and de-capping (Coller & Parker, 2004; Di Giammartino, Nishida, & Manley, 2011). This layer of regulation contributes to the amount of protein produced per transcript. Failure to properly process the pre-mRNA in the nucleus or the mature mRNA in the cytoplasm can lead to dysregulation of protein production and disease. Post-transcriptional regulation is especially critical in developmental processes such as gametogenesis and embryogenesis (Salles, Lieberfarb, Wreden, Gergen, & Strickland, 1994; Y. Zhang, Park, Blaser, & Sheets, 2014). During the early stages of embryogenesis prior to the onset of zygotic transcription, inherited maternal mRNAs and proteins are critical to axis formation and cell fate specification (Bashirullah et al., 1999; Tao et al., 2005; J. Zhang et al., 1998). Maternal mRNAs must be produced in the germline, packaged into oocytes, silenced, activated at the right time and place in the embryo, and then cleared once zygotic transcription begins. Accordingly, a variety of post-transcriptional regulatory mechanisms are required to coordinate this developmental program. Much remains to be learned about how they collaborate to achieve distinct spatiotemporal expression patterns for different maternal mRNAs.

The germline of the hermaphroditic nematode *Caenorhabditis elegans* is a suitable model for studying spatiotemporal regulation of maternal mRNA (Hubbard & Greenstein, 2005; Lee & Schedl, 2006). The gonads consist of two symmetrical tube-shaped tissues that contain mitotically dividing germ cells in the distal end of each tube (Fig. 1a). As the mitotic nuclei move away from the distal end, they begin to enter meiosis I and form a syncytium where the nuclei migrate to the periphery and share cytoplasmic content. The nuclei recellularize to form oocytes near the loop region as the tube bends. In the proximal end, oogenesis is followed by fertilization in the spermatheca where the sperm produced during the L4 larval stage or acquired by mating is stored. Embryogenesis continues in the uterus, where a hard chitin shell is secreted (Strome et al., 1994). The 1-cell embryo undergoes multiple pre-ordained cellular divisions to establish the body axes, segregate germline from soma, and define the number of tissue lineages before it exits the uterus (Hubbard & Greenstein, 2005).

**Figure 1.**
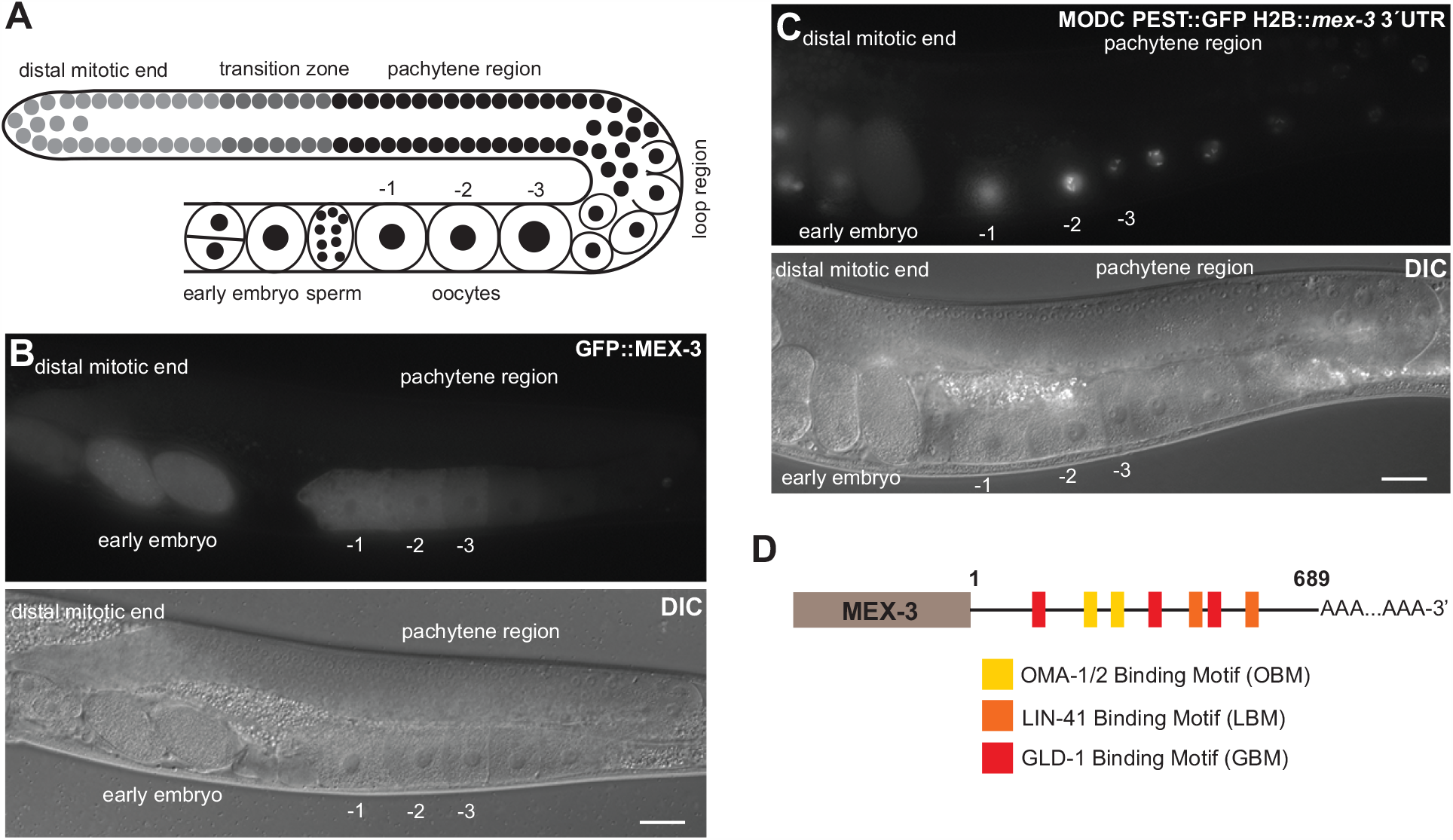
MEX-3 exhibits a unique expression pattern in the germline. **(A)** A schematic representing germline organization in *C. elegans*. One of two gonadal arms is shown. Germ cells undergo mitotic divisions in the distal end and then enter meiosis as they move farther from the distal tip cell. The syncytial meiotic nuclei start to recellularize around the loop region to form oocytes. In the proximal end, late oocytes undergo maturation, get fertilized by the sperm, and then move to the uterus to undergo embryonic development. **(B)** DIC and fluorescence images of an adult hermaphrodite germline from the strain in which MEX-3 is endogenously tagged with GFP (GFP::MEX-3). MEX-3 is present in the distal mitotic end, maturing oocytes, and early embryo. **(C)** DIC and fluorescence images of an adult hermaphrodite germline from the transgenic reporter strain carrying a germline promoter fused to GFP and the *mex-3* 3’UTR. MEX-3 is present in the distal mitotic end, maturing oocytes, and early embryo. **(D)** A schematic representing the 3’UTR of *mex-3* and some of its putative binding motifs. Images taken at 40x magnification. Scale bars = 30µm.

Post-transcriptional regulatory mechanisms contribute to each of these processes in the germline. For instance, the conserved maxi KH-domain RNA-binding protein GLD-1 promotes entry into meiosis in part by binding a specific regulatory element in the 3’UTR of the mitosis-promoting notch receptor *glp-1* and repressing its translation (Marin & Evans, 2003), the PUF-domain RNA-binding proteins FBF-1/2 promote mitosis in the distal end through 3’UTR-mediated translational repression of *gld-1* (Suh et al., 2009), and zinc finger RNA-binding proteins OMA-1/2 promote oocyte maturation in the proximal end through repression of multiple transcripts (Guven-Ozkan, Robertson, Nishi, & Lin, 2010; Kaymak & Ryder, 2013; Lin, 2003; Spike et al., 2014).

The highly conserved KH-domain RNA-binding protein MEX-3 promotes anterior cell fate specification in the embryo and contributes to maintenance of totipotency in the germline (Ciosk, DePalma, & Priess, 2006; Draper, Mello, Bowerman, Hardin, & Priess, 1996; Huang & Hunter, 2015). Null mutants of *mex-3* are maternal-effect embryonic lethal where the embryos fail to hatch due to cell fate patterning defects (Draper et al., 1996). MEX-3 is evolutionarily conserved across multicellular animals. There are four human MEX-3 homologues (hMEX-3A-D) (Pereira, Le Borgne, Chartier, Billaud, & Almeida, 2013). Some of these proteins function in cellular differentiation pathways (Buchet-Poyau et al., 2007; Pereira, Sousa, et al., 2013). For example, hMEX-3A regulates intestinal cell fate specification by 3’UTR-mediated negative regulation of the *cdx2* mRNA, which encodes a homeobox transcriptional factor (Pereira, Sousa, et al., 2013). hMEX-3A expression is upregulated in gastric cancer (Jiang et al., 2012). The planarian homologue (mex3-1) maintains the pool of mitotic stem cells in addition to promoting stem cell differentiation (Zhu, Hallows, Currie, Xu, & Pearson, 2015). In the fish *Nothobranchius furzeri*, mex3A contributes to maintenance of proliferating neuronal stem cells (Naef et al., 2020).

*C. elegans* MEX-3 binds two short motifs separated by zero to eight bases ((A/G/U)(G/U)AGN_(0–8)_U(U/A/C)UA) (Pagano, Farley, Essien, & Ryder, 2009). MEX-3 is present in the distal mitotic end, maturing oocytes, and the early embryo where it also associates with P-granules (Draper et al., 1996)--membrane-less structures composed of RNA and protein. MEX-3 contributes to establishing the anterior/posterior asymmetry in the 1-cell embryo by repressing *pal-1* mRNA in the anterior blastomere and therefore restricting it to the posterior blastomere where PAL-1 is necessary for posterior cell fate specification (Huang & Hunter, 2015; Huang, Mootz, Walhout, Vidal, & Hunter, 2002). MEX-3 also plays a role in maintaining totipotency in the germline; animals carrying null mutations in both *gld-1* and *mex-3* exhibit signs of transdifferentiation of the germ cells to neuronal or pharyngeal cells (Ciosk et al., 2006). Although we know some of the downstream target mRNAs of MEX-3, we do not know how *mex-3* mRNA itself is regulated. We previously discovered that the *mex-3* 3’UTR is sufficient to confer the MEX-3 pattern of expression to a reporter gene. Transgenic animals carrying a reporter transgene driven by a pan-germline promoter fused to GFP and the 3’UTR of *mex-3* ((*Pmex-5::MODC PEST::GFP::H2B:: mex-3 3*’*UTR)*, MODC: Mouse Ornithine DeCarboxylase) exhibit an expression pattern that is similar to that of the endogenous MEX-3 (Kaymak et al., 2016) (Fig. 1). Additionally, the 3’UTR of *mex-3* contains putative binding motifs for several germline RBPs such as GLD-1, LIN-41, and OMA-1/2, but it remains unknown whether these binding motifs are functional and if they contribute to the spatiotemporal localization of MEX-3 and animal fertility.

In this study, we demonstrate that *mex-3* mRNA is indeed post-transcriptionally regulated. We identify the RNA-binding proteins that control its germline spatiotemporal expression through its 3’UTR, as well as the regulatory mechanisms involved. We find that different mechanisms govern the expression pattern of MEX-3 in different regions of the germline, leading to its differential abundance across the gonad. We also show that the 3’UTR of *mex-3* is surprisingly dispensable for fertility but does contribute to animal fecundity. Overall, our data define a model for how MEX-3 is patterned and demonstrate that the primary role of its 3’UTR is to enhance reproductive robustness.

## Results

### DAZ-1, GLD-1, LIN-41, and OMA-1/2 regulate MEX-3 expression in the germline

MEX-3 exhibits a unique expression pattern (Fig.1b). In wild type animals, MEX-3 is expressed at low levels in the distal mitotic progenitor cells and is absent in the syncytial meiotic region of the germline. MEX-3 is also expressed in early immature oocytes with its overall abundance progressively increasing as the oocytes approach maturation (Fig. 1b). Transcripts encoding *mex-3* have a long 3’UTR (689bp) that contains putative binding motifs for several germline RBPs (Fig. 1d). We previously demonstrated that this 3’UTR is sufficient to confer the MEX-3 pattern of expression to a reporter gene in live animals (Fig. 1c) (Kaymak et al., 2016). Thus, we predicted that the germline RBPs that associate with these motifs might coordinate MEX-3 expression through its 3’UTR via post-transcriptional regulatory mechanisms.

To test this hypothesis, we performed an RBP-targeted RNAi screen using a strain in which wild type MEX-3 is endogenously tagged with GFP (GFP::MEX-3) (Tsukamoto et al., 2017). We soaked L4 or arrested L1 larval stage animals in dsRNA corresponding to the coding sequence of the candidate RBP. Then, we placed the animals on *E. coli* OP50 as a food source and imaged the adults using fluorescence microscopy to assess the effect of the RNAi on the germline GFP::MEX-3 expression pattern. We found that knockdown of *daz-1, gld-1, lin-41*, or *oma-1/2* significantly altered the pattern of wild type GFP::MEX-3 expression (Fig. 2, Table S4). Knockdown of *daz-1* resulted in an overall increase of GFP::MEX-3 in the distal mitotic region (Fig. 2b, 2f, Table S4), but had no impact on the syncytial meiotic pachytene region.

**Figure 2.**
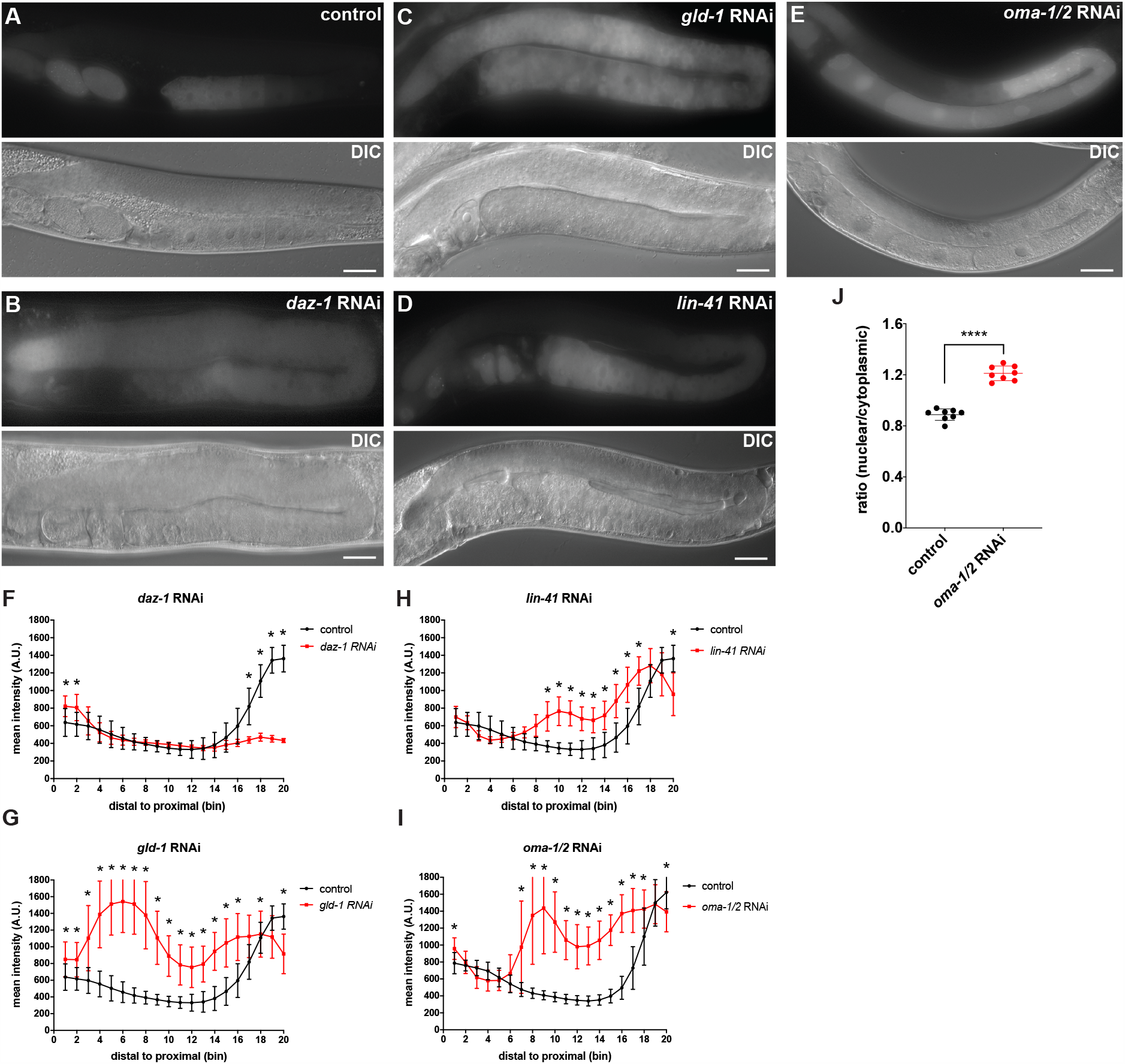
DAZ-1, GLD-1, OMA-1/2, and LIN-41 regulate spatiotemporal expression pattern of MEX-3 in the germline. **(A)** DIC and fluorescence images of wild type GFP::MEX-3 animals from the control RNAi. **(B)** DIC and fluorescence images of GFP::MEX-3 animals after *daz-1* knockdown. GFP::MEX-3 was significantly increased in the mitotic distal end. **(C)** DIC and fluorescence images of GFP::MEX-3 animals after *gld-1* knockdown. GFP::MEX-3 expression was derepressed in the meiotic region. **(D)** DIC and fluorescence images of GFP::MEX-3 animals after *lin-41* knockdown. GFP::MEX-3 was de-repressed in the loop region. **(E)** DIC and fluorescence images of GFP::MEX-3 animals after *oma-1/2* knockdown. GFP-MEX-3 was significantly increased in the oocytes. **(F)** quantitative analysis of fluorescence intensity after *daz-1* knockdown (n=9/15). For all the images from the RNAi, a line with a width of 30 pixels was drawn along the entire germline and fluorescence intensities were binned (20 bins). Data are shown as the mean fluorescence intensity ± standard deviation (SD). A two tailed student t-test was performed to compare the means for each bin from control animals and the RNAi condition to assess significance. For RNAi conditions that have the same control, a one-way unstacked ANOVA was used to assess the overall significance, then Bonferroni adjusted p-values were calculated by multiplying pairwise Fisher’s LSD test p-values by the number of hypotheses tested. All p-values for this figure are reported in table S4. **(G)** quantitative analysis of fluorescence intensity after *gld-1* knockdown (n=7/13). **(H)** quantitative analysis of fluorescence intensity after *lin-41* knockdown (n=17/17). **(I)** quantitative analysis of fluorescence intensity after *oma-1/2* knockdown (n=9/9). **(J)** quantitative analysis of nuclear GFP::MEX-3 of the *oma-1/2* RNAi animals. Nuclear fluorescence intensity was divided by the cytoplasmic fluorescence intensity for each oocyte. Each dot represents the averaged ratios from the two most proximal oocytes in an individual animal. (*) indicates statistical significance, adjusted p-value ≤ 0.05. (****) indicates statistical significance, p-value ≤ 0.0005. Scale bar = 30 µm.

Knockdown of *daz-1* also resulted in defective oogenesis, so the impact on GFP::MEX-3 in the oocytes could not be directly assessed but appeared reduced in the quantitation due to the absence of oocytes. Knockdown of *gld-1* resulted in increased expression of GFP::MEX-3 in the distal mitotic end as well as expansion of GFP::MEX-3 to the meiotic region (Fig. 2c, 2g, Table S4). These results confirm a previous finding using anti-MEX-3 antibody staining after *gld-1* RNAi (Mootz, Ho, & Hunter, 2004). Knockdown of *lin-41* resulted in expansion of GFP::MEX-3 to the loop region (Fig. 2d, 2h, Table S4), while knockdown of *oma-1/2* caused an increase in GFP::MEX-3 expression in the oocytes, also confirming prior observations (Tsukamoto et al., 2017). We also note that knockdown of *oma-1/2* led to accumulation of GFP::MEX-3 in the nucleus (Fig. 2e, 2i, 2j, Table S4). By contrast, knockdown of *pos-1* or *pie-1* had no effect on the pattern of expression. Together, our data identify DAZ-1 as a regulator of GFP::MEX-3 expression in the distal mitotic end and confirm that GLD-1, LIN-41, and OMA-1/2 regulate GFP::MEX-3 expression in the pachytene region, the loop region, and in late oocytes, respectively.

To determine whether the RBPs identified above regulate *mex-3* expression through its 3’UTR, we knocked down *daz-1, gld-1, lin-41*, or *oma-1/2* in the transgenic reporter strain described above where a pan-germline promoter (*Pmex-5*) drives the expression of a nuclear MODC PEST::GFP*::H2B* reporter under the control of the *mex-3* 3’UTR. Knockdown of *gld-1, lin-41*, or *oma-1/2* changed the pattern of reporter expression similarly to knockdown of endogenous GFP::MEX-3 (Fig. S1, Table S7), suggesting that these RBPs act through the *mex-3* 3’UTR. By contrast, knockdown of *daz-1* did not show a strong increase of reporter expression in the distal mitotic end, suggesting that this protein alters MEX-3 expression via a 3’UTR-independent mechanism, possibly through the coding sequence, the 5’end, or indirectly through dysregulation of factors that act on the *mex-3* promoter.

### Poly(A) tail length control mediates the spatiotemporal expression pattern of MEX-3

The length of the poly-adenosine tail contributes to the stability and translational efficiency of eukaryotic mRNAs and has been demonstrated to control post-transcriptional regulation of maternal mRNAs in several metazoans including *C. elegans* (Lima et al., 2017; Nousch, Yeroslaviz, Habermann, & Eckmann, 2014; Salles et al., 1994). To test whether cytoplasmic polyadenylation contributes to the pattern of MEX-3 expression, we used RNAi to knock down components of the germline cytoplasmic poly(A) polymerase complexes (*gld-2, gld-4, gld-3, rnp-8*) in the wild type GFP::MEX-3 strain (Fig. 3, Table S5). Knockdown of *gld-2* resulted in reduced GFP::MEX-3 expression in the transition zone/early meiotic region and the immature oocytes (Fig. 3b, 3g, Table S5). Since GLD-2 does not contain an RNA-binding domain, it requires an RBP co-factor such as GLD-3 or RNP-8 to direct it to specific germline transcripts (Eckmann, Crittenden, Suh, & Kimble, 2004). Knockdown of *gld-3* resulted in increased GFP::MEX-3 expression throughout the oogenic region (Fig. 3c, 3h, Table S5). In treated animals, GFP::MEX-3 also accumulated in punctate-like formations surrounding the nucleus and near the plasma membrane. GLD-3 promotes spermatogenesis in the sex-determination pathway (Eckmann, Kraemer, Wickens, & Kimble, 2002). Thus, knockdown of *gld-3* results in spermatogenesis defects, causing oocytes to stack in the proximal germline and expand to the loop (Fig. 3c, 3h). As such, the enhanced GFP::MEX-3 fluorescence could be the result of an accumulation of mature oocytes. It could also be that GLD-3, independently of GLD-2, represses MEX-3 through an unknown pathway. Neither knockdown of *gld-4* nor *rnp-8* changed the expression of GFP::MEX-3. Together, these data suggest that GLD-2 and GLD-3 play a role in regulating wild type GFP::MEX-3 expression in the germline, but given their differential effect, it is likely that they do not act in the same way.

**Figure 3.**
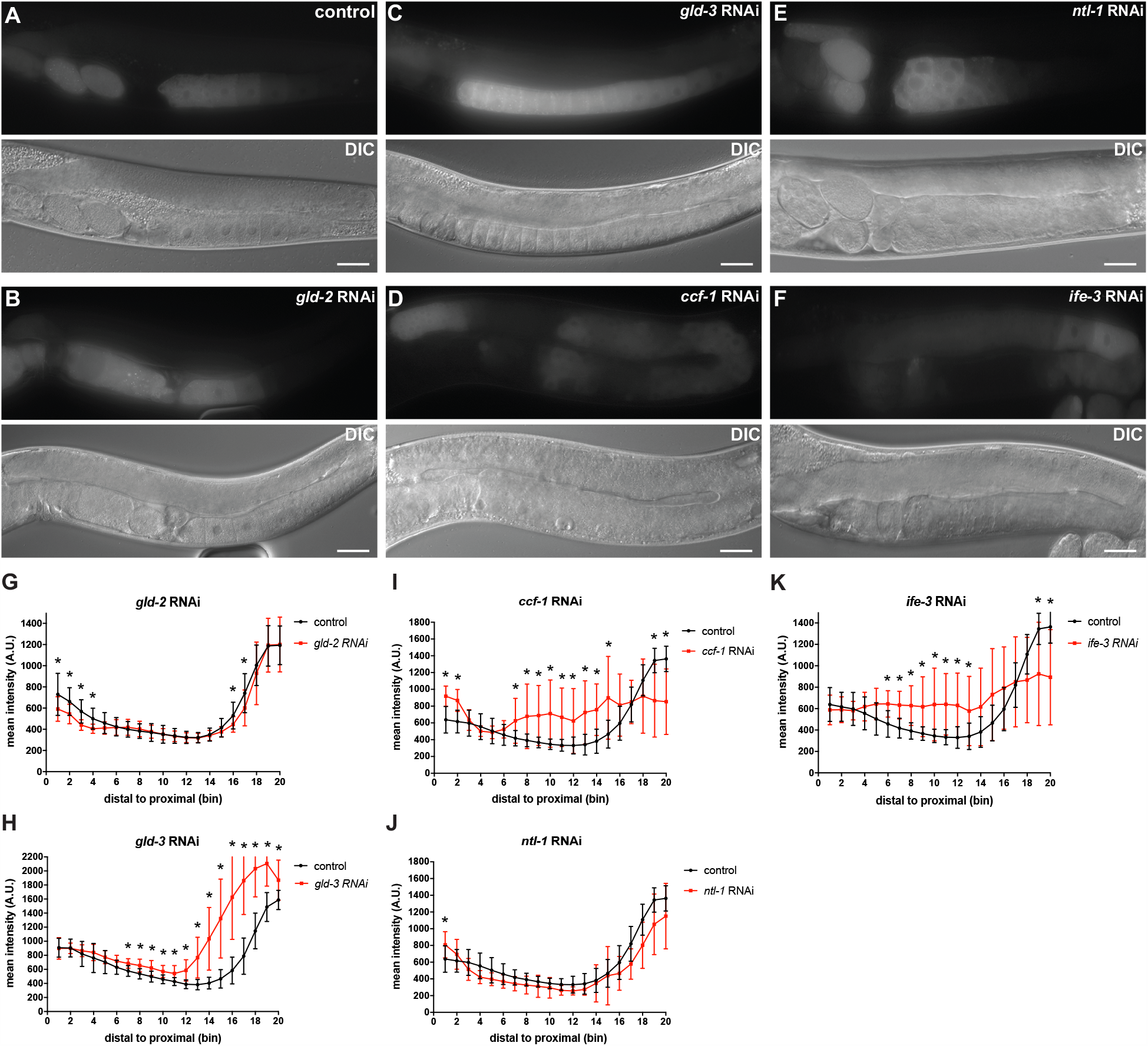
Poly(A) tail length control and translational regulation contribute to post-transcriptional regulation of *mex-3* in the germline. **(A)** DIC and fluorescence images of wild type GFP::MEX-3 animals from the control RNAi. **(B)** DIC and fluorescence images after *gld-2* knockdown. GFP::MEX-3 was significantly reduced in the distal mitotic end and oocytes. **(C)** DIC and fluorescence images after *gld-3* knockdown. GFP::MEX-3 expression was significantly increased in the oogenic region. **(D)** DIC and fluorescence images after *ccf-1* knockdown. GFP::MEX-3 expression was significantly increased in the distal mitotic end and the meiotic region. **(E)** DIC and fluorescence images after *ntl-1* knockdown. GFP::MEX-3 expression was significantly increased in the distal mitotic end. **(F)** DIC and fluorescence images after *ife-3* knockdown. GFP::MEX-3 expression was significantly increased in the distal end and meiotic region. **(G)** quantitative analysis of fluorescence intensity after *gld-2* knockdown (n=15/15). **(H)** quantitative analysis of fluorescence intensity after *gld-3* knockdown (n=7/12). **(I)** quantitative analysis of fluorescence intensity after *ccf-1* knockdown (n=15/15). **(J)** quantitative analysis of fluorescence intensity after *ntl-1* knockdown (n=13/13). **(K)** quantitative analysis of fluorescence intensity after *ife-3* knockdown (n=13/13). (*) indicates statistical significance, adjusted p-value ≤ 0.05. All p-values for this figure are reported in table S5. Scale bar = 30 µm.

In *C. elegans*, the major deadenylation complex consists of the subunits CCF-1, CCR-4, and NTL-1. To assess the role of cytoplasmic deadenylation in repressing GFP::MEX-3, we knocked down these components using RNAi. Knockdown of either *ccf-1* or *ntl-1* altered the expression pattern of GFP::MEX-3 (Fig. 3d-3e, 3i-3j, Table S5). Knockdown of *ccf-1* causes meiotic defects that lead to formation of small cellularized nuclei that look like oocytes but the nuclei are arrested in pachytene (Molin & Puisieux, 2005). Knockdown of *ccf-1* resulted in increased expression in the mitotic region, expansion to early meiotic zone, and ectopic expression in the oocytes where some defective oocytes appeared to have varying levels of GFP::MEX-3 (Fig. 3d, 3i, Table S5). Knockdown of *ntl-1* causes defects in meiotic progression, preventing formation of normal oocytes and leading to defects in germline organization where small defective oocytes appear in multiple layers in the proximal end. Knockdown of *ntl-1* resulted in increased expression of GFP::MEX-3 in the mitotic region (Fig. 3e, 3j, Table S5). Knockdown of *ccr-4* did not alter the GFP::MEX-3 expression suggesting that CCR-4 alone is not essential for regulating MEX-3 expression or that it is not a key component of the deadenylation complex that regulates MEX-3 expression. Together, these results show that components of the cytoplasmic polyadenylation machinery positively regulate MEX-3 expression in the mitotic and oogenic regions, while components of the deadenylation machinery repress GFP::MEX-3 expression in the mitotic, meiotic, and oogenic regions. We propose that differential activity of each pathway, in different regions of the germline, coordinates the overall pattern of endogenous MEX-3 expression.

### The translation initiation factor IFE-3 represses the expression of MEX-3 in the mitotic and meiotic regions

Among the five *C. elegans* translation initiation factor eIF4E homologs, only *ife-3* causes embryonic lethality when knocked down (Keiper et al., 2000). A recent report revealed that this factor negatively regulates the translation of specific maternal transcripts in the germline, presumably by interfering with normal translation initiation mediated by the other homologs (Huggins et al., 2020). To test whether *ife-3* contributes to the pattern of MEX-3 expression, we knocked it down via RNAi in the wild type GFP::MEX-3 strain. Knockdown of *ife-3* caused defects in late stages of meiosis leading to the failure to form oocytes (Fig. 3f, 3k, Table S5). In addition, we observed an increase of GFP::MEX-3 levels in the meiotic region. These results suggest that IFE-3 may be involved in mechanisms that regulate translation and expression of MEX-3 in the meiotic region.

### The 3’UTR of *mex-3* is required for the spatiotemporal expression of MEX-3

The 3’UTR of *mex-3* contains putative binding motifs for several germline RNA-binding proteins, including those that we and others have shown contribute to its pattern of expression above (Fig. 1d, Fig. 2). To investigate whether these binding motifs contribute to the pattern of MEX-3 expression pattern, we used CRISPR/Cas9 to make an allelic series of *mex-3* 3’UTR deletion mutants in the wild type GFP::MEX-3 background (Fig. 4a), enabling observation of changes in the expression pattern of GFP::MEX-3 in addition to scoring the resulting phenotypes. We generated three small deletions (*spr6*: 142bp, *spr10*: 190bp, *spr7*: 134bp) and two larger deletions (*spr5*: 488bp, *spr9*: 624bp) in this series (Fig. 4a). All mutants were made in the wild type GFP::MEX-3 background except *mex-3(spr5). mex-3(spr5)* was made in a background strain in which wild type MEX-3 is not tagged with GFP. Among the four mutants made in the GFP::MEX-3 background, only *mex-3(spr9)* and *mex-3(spr10)* mutant animals exhibited altered expression pattern. *mex-3(spr9)* mutant animals exhibited a significant increase in GFP::MEX-3 expression throughout the germline, especially in the meiotic and loop regions (Fig. 4b, 4c, Table S6). *mex-3(spr10)* mutant animals showed a modest increase of GFP::MEX-3 in the loop region (Fig. 4d, 4e, Table S6).

**Figure 4.**
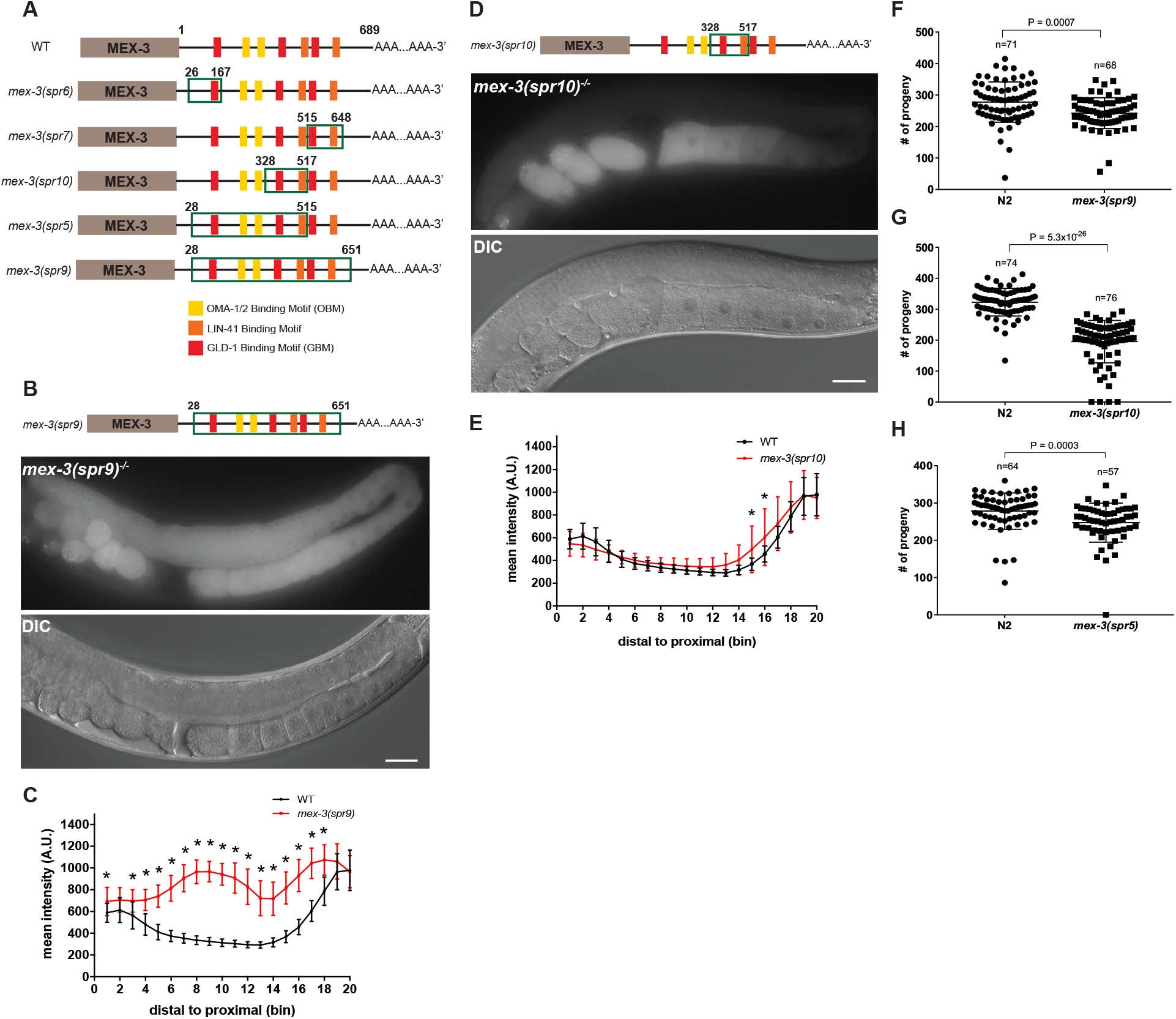
*mex-3* 3’UTR deletions alter MEX-3 expression and reduce fertility. **(A)** A schematic representing the 3’UTR deletions made in the wild type GFP::MEX-3 strain. The region deleted in each mutant is highlighted by the green rectangle. **(B)** DIC and fluorescence images of the germline of the *mex-3(spr9)* homozygous mutant animals. **(C)** quantitative analysis of the fluorescence intensity in the *mex-3(spr9)* mutant animals compared to that of wild type GFP::MEX-3 (n=23). **(D)** DIC and fluorescence images of the germline of the *mex-3(spr10)* homozygous mutant animals. **(E)** quantitative analysis of the fluorescence intensity in the *mex-3(spr10)* mutant animals compared to that of wild type GFP::MEX-3 (n=17). **(F)** brood size assay of *mex-3(spr9)* mutant animals at 20°C. Each dot represents the brood size of an individual animal. Data from three biological replicates are shown in the graph. P-values are from a Kolmogorov-Smirnov test. **(G)** brood size assay of *mex-3(spr10)* mutant animals at 20°C. **(H)** brood size assay of *mex-3(spr5)* mutant animals at 20°C. All p-values for panels C and E are reported in table S6. All images taken at 40x magnification. Scale bar = 30 µm.

The altered expression pattern of GFP::MEX-3 throughout the germline caused by deleting majority of the 3’UTR (*spr9*) indicates that the 3’UTR contains various cis-regulatory elements that coordinate GFP::MEX-3 expression through multiple mechanisms in different regions of the germline. The de-repression of GFP::MEX-3 observed in the loop region in the *mex-3(spr10)* mutant animals indicates that the deletion in this mutant may contain repressive elements that mediate repression of *mex-3* in that region. To assess whether any of the 3’UTR deletions disrupt poly(A) processing leading to aberrant 3’-end formation, we used a poly(A) tail-driven approach to amplify and sequence the 3’-end of *mex-3* transcripts produced by each mutant. None of the deletions affected the poly(A) processing site selection. All mutants use the most common poly(A) processing site found in endogenous *mex-3* (Table S8).

### The 3’UTR of *mex-3* is not required for viability but contributes to animal fecundity

All five mutants including the *spr9* allele that deletes majority of the 3’UTR (624bp) are viable as homozygotes and can be easily propagated as such. This demonstrates that the 3’UTR, though sufficient to pattern reporter expression, is not essential for viability. To determine if any of the mutations compromise reproductive health, we measured the brood size in animals homozygous for the deletions compared to control wild type animals. Three of the five deletion mutants exhibited reduced fertility. Brood size was reduced in *mex-3(spr9)* (Fig. 4f, p-value = 0.0007), *mex-3(spr10)* (Fig. 4g, p-value = 5.3*10^−26^), and *mex-3(spr5)* (Fig. 4h, p-value = 0.0003).

Moreover, *mex-3(spr9)* and *mex-3(spr10)* mutant animals exhibited partial sterility at 25°C (Fig. 5). 2.5% of homozygous *mex-3(spr9)* mutant animals were sterile while 6.25% of the homozygous *mex-3(spr10)* mutant animals were sterile (Fig. 5C). Sterile animals from both strains exhibited a GFP::MEX-3 expression pattern similar to that of the non-sterile homozygous mutant animals. However, sterile animals from both strains exhibited defects in oogenesis (Fig. 5a-5b). The gonads did not contain normal oocytes or viable embryos. The reduction in brood size, partial sterility, and germline defects in the sterile animals indicate that the 3’UTR of *mex-3* contributes to animal fecundity through ensuring normal gametogenesis.

**Figure 5.**
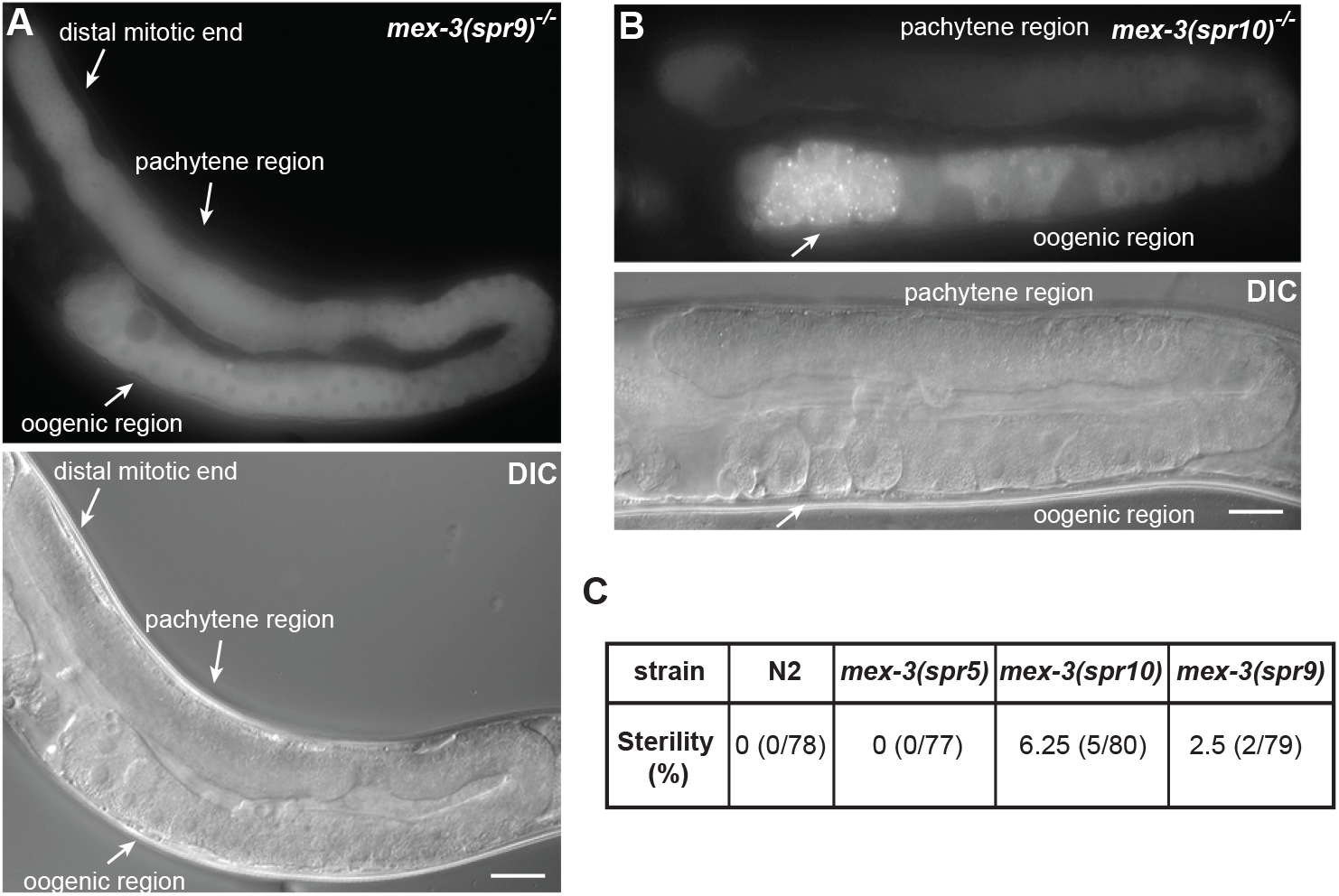
*mex-3(spr9)* and *mex-3(spr10)* mutant animals exhibit partial sterility. **(A)** a representative image of a sterile *mex-3(spr9)* homozygous mutant animal at 25°C. The animals fail to produce normal oocytes or viable embryos. Oogenesis appears to be defective. **(B)** a representative image of a sterile *mex-3(spr10)* homozygous mutant animal at 25°C. The animals contain smaller than normal defective oocytes. GFP::MEX-3 appears to accumulate in granules. **(C)** a table showing the percentage of sterile animals in each mutant population as well as the wild type animals. Each adult was grown at 25°C and its progeny scored for fertility/sterility. All images taken at 40x magnification. Scale bar = 30 µm.

## Discussion

### DAZ-1 regulation of MEX-3 expression in the mitotic progenitor cells

The 3’UTR of maternal germline mRNAs contain binding motifs for numerous germline RNA-binding proteins (Aeschimann et al., 2017; Farley, Pagano, & Ryder, 2008; Kaymak & Ryder, 2013; Ryder, Frater, Abramovitz, Goodwin, & Williamson, 2004; Tamburino, Ryder, & Walhout, 2013). However, we don’t know which of the predicted motifs are functional and whether these 3’UTRs are required for germline development. Here, we dissected the 3’UTR of the *mex-3* gene, which encodes an RNA-binding protein required for embryonic cell fate patterning and totipotency in the germline. MEX-3 is expressed in the distal mitotic end, maturing oocytes, and the early embryo (Draper et al., 1996; Tsukamoto et al., 2017) (Fig. 1b, 1c). Additionally, the 3’UTR of *mex-3* contains putative binding motifs for several germline RNA-binding proteins (Fig. 1d). Our candidate RBP RNAi screen revealed that knockdown of *daz-1, gld-1, lin-41*, or *oma-1/2* altered GFP-tagged endogenous MEX-3 (GFP::MEX-3) expression in differing region in the germline (Fig. 2)

DAZ-1, an RNA-binding protein that contains an RRM (RNA Recognition Motif) and contributes to meiotic progression during oocyte development (Karashima, Sugimoto, & Yamamoto, 2000), appears to regulate the expression pattern of MEX-3 in the mitotic region. Although MEX-3 is expressed in that region in wild type animals, knockdown of *daz-1* caused a significant increase of GFP::MEX-3 expression in the distal mitotic end (Fig. 2b, 2f, Table S4), suggesting that DAZ-1 represses *mex-3* in that region. The binding specificity of DAZ-1 is unknown. Therefore, we don’t know if the 3’UTR of *mex-3* contains binding motifs for DAZ-1. However, knockdown of *daz-1* in the *mex-3* 3’UTR reporter strain did not show similar results in the distal mitotic end (Fig. S1). Therefore, DAZ-1 likely does not regulate *mex-3* expression through its 3’UTR. Interestingly, GFP::MEX-3 was significantly increased in the *mex-3(spr9)* mutant animals in the distal mitotic end (Fig. 4b, 4c, Table S6) indicating that the 3’UTR contains cis-regulatory elements that negatively regulate its expression in the mitotic progenitor cells. Although DAZ-1 may not regulate MEX-3 expression through its 3’UTR, it is possible that it may regulate the expression of other pathways that directly influence *mex-3* expression (Fig. 6a). The increased MEX-3 expression in the distal mitotic end as a result of knockdown of the deadenylation complex components *ccf-1* or *ntl-1* is consistent with this possibility. The presence of a mechanism to repress MEX-3 expression in the distal mitotic end suggests that overexpression of MEX-3 in the distal end may negatively impact mitotically dividing germ cells, in certain contexts. MEX-3 has been shown to contribute to maintenance of totipotency by repressing *pal-1*, which promotes development of body muscles (Ciosk et al., 2006). Precisely how DAZ-1 contributes to this pattern, and whether it works through cytoplasmic deadenylation, is still unknown.

**Figure 6.**
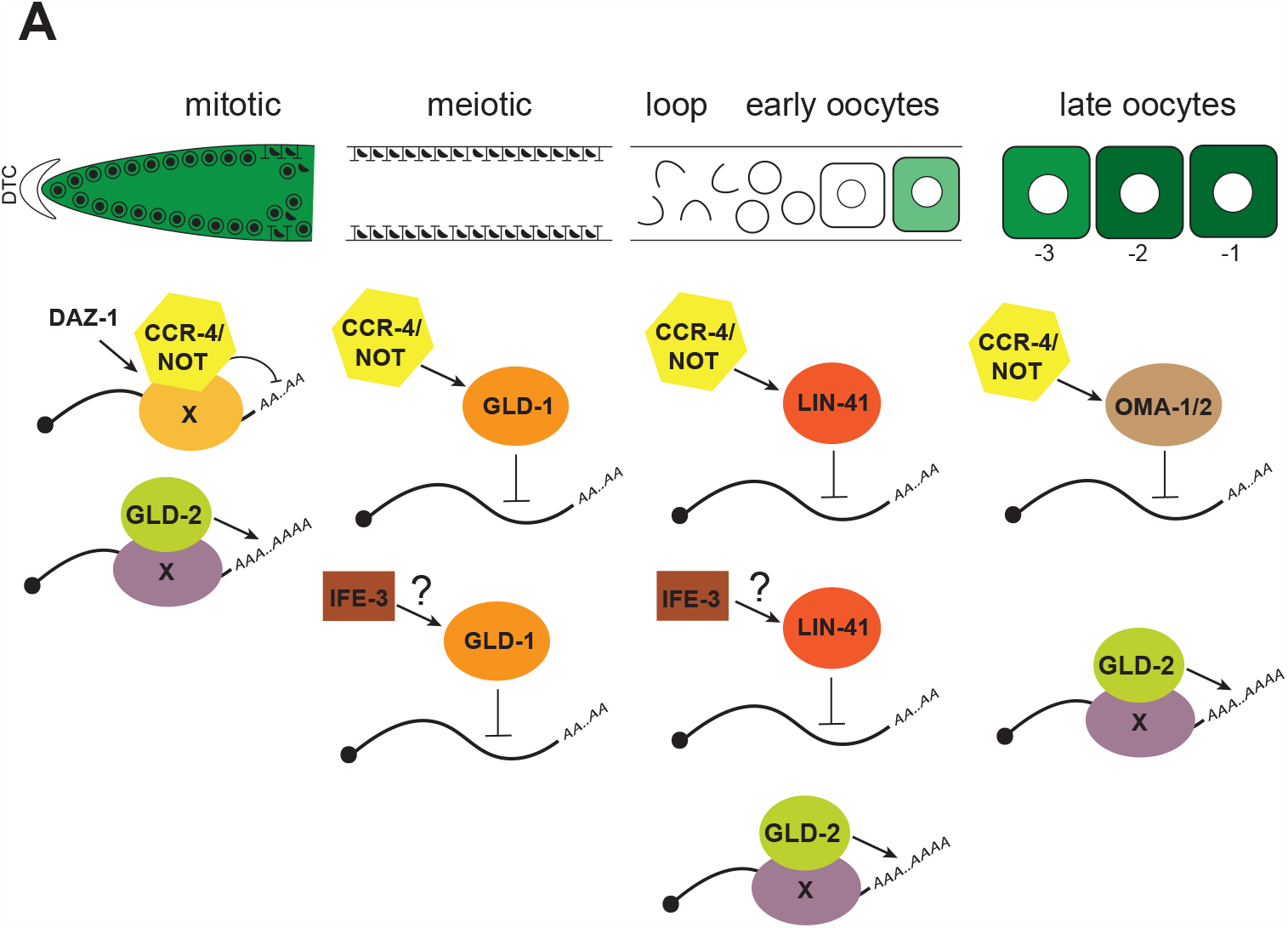
Model for 3’UTR-mediated post-transcriptional regulation of *mex-3* in the germline. **(A)** DAZ-1 indirectly regulates expression of MEX-3 in the distal mitotic end. GLD-1 represses MEX-3 expression through its 3’UTR in the meiotic region. LIN-41 represses expression of MEX-3 in the loop region while OMA-1/2 repress expression of MEX-3 in the maturing oocytes. Cytoplasmic polyadenylation positively regulates expression of MEX-3 in the distal mitotic end and oocytes while cytoplasmic deadenylation negatively regulates expression of MEX-3 throughout the entire germline. IFE-3 negatively regulates expression of MEX-3 in the meiotic and loop regions.

### GLD-1-mediated repression of endogenous MEX-3 expression in the meiotic region

Our results are consistent with and expand upon a previous study that showed GLD-1 represses *mex-3* expression in the germline (Mootz et al., 2004). The 3’UTR of *mex-3* also contains three putative GLD-1 binding motifs (GBMs) (Ryder et al., 2004; Wright et al., 2011). Consistent with previous findings, our results reveal that *gld-1* knockdown leads to de-repression of the endogenous GFP::MEX-3 in the meiotic region (Fig. 2c, 2g, Table S4). The observation of a similar result in the *mex-3* 3’UTR reporter transgenic strain (Fig. S1, Table S7) indicates that this repression acts through the 3’UTR. Among the 3’UTR deletion mutants, the *mex-3(spr7)* deletion removes a single GBM but displays no altered expression in the meiotic region (Fig. S2, Table S6). The same is also true in the *mex-3(spr6)* mutant animals, where the deletion removes a different GBM (Fig. S2, Table S6) and *mex-3(spr10)* mutant animals where the 3’UTR deletion removes the third GBM (Fig. 4d, 4e, Table S6). These results indicate that loss of any one of the three putative GBMs is not sufficient to de-repress *mex-3* expression on its own. However, *mex-3(spr9)* mutant animals in which all three GBMs are deleted show complete de-repression of MEX-3 in the meiotic region (Fig. 4b, 4c, Table S6). It is also possible that other RBPs contribute to the 3’UTR-mediated repression of *mex-3* in the meiotic region. The deletion in the *mex-3(spr5)* mutant animals removes two GBMs, but we do not know how the deletion impacts the expression of endogenous MEX-3. GLD-1 may repress *mex-3* expression by binding the deadenylation complex and promoting deadenylation of *mex-3* mRNA reducing its stability and leading to its degradation. GLD-1 may also repress *mex-3* expression by inhibiting its translation (Lee & Schedl, 2004). Consistent with this hypothesis, knockdown of deadenylation components *ccf-1* and *ntl-1*, or knockdown of the translational repressor *ife-3*, partially phenocopies the *spr9* mutant where all three GBMs are deleted (Fig. 3d-3f, Fig. 4b). Taken together, our findings support a model where GLD-1 represses *mex-3* translation directly through its 3’UTR (Fig. 6a), and that multiple binding motifs can contribute to repression.

### LIN-41-mediated repression of endogenous MEX-3 expression in the loop region

LIN-41 is expressed in the loop region, where meiotic nuclei start to recellularize, and in early immature oocytes (Tsukamoto et al., 2017). The 3’UTR of *mex-3* contains two putative LIN-41 binding motifs (LBM). GFP::MEX-3 expression was previously shown to expand to the loop region in a *lin-41* null mutant background (Tsukamoto et al., 2017). Consistent with these findings, we observed expansion of GFP::MEX-3 to the loop region when we knocked down *lin-41* in the wild type GFP::MEX-3 strain (Fig. 2d, 2h, Table S4). We observed similar results when we knocked down *lin-41* in the *mex-3* 3’UTR reporter strain (Fig. S1, Table S7), suggesting that LIN-41 represses *mex-3* in the loop region through its 3’UTR. This is also consistent with a study that showed *mex-3* mRNA associates with purified recombinant LIN-41 in an *in vitro* pull-down assay (Tsukamoto et al., 2017). Additionally, the 3’UTR deletion in the *mex-3(spr7)* mutant animals removes one putative LIN-41 binding motif while the 3’UTR deletion in the *mex-3(spr10)* mutant animals removes the other putative LIN-41 binding motif (Fig. 4a). The *mex-3(spr10)* mutant animals showed de-repression of MEX-3 in the loop region while *mex-3(spr7)* mutant animals did not, suggesting that the motif deleted in the *spr10* allele is partially sufficient to repress *mex-3* in the loop region (Fig. S2, Fig. 4d, 4e). Interestingly, *mex-3(spr9)* mutant animals which delete both putative binding motifs showed strong de-repression of MEX-3 in the loop region (Fig. 4b, 4c). This observation suggests that both motifs may be required for complete repression of MEX-3 in the loop region. Alternatively, given that the deletion in the *mex-3(spr9)* mutant animals deletes most of the 3’UTR, it could also be that there are additional RBPs that mediate repression of MEX-3 through its 3’UTR in that region. Together, our findings show that the 3’UTR of *mex-3* is required for repression of MEX-3 in the loop region (Fig. 6a), and that one of two LIN-41 motifs is sufficient to partially repress MEX-3 expression in this region.

### OMA-1/2-mediated repression of endogenous MEX-3 expression in the oocytes

OMA-1/2 are expressed in the oocytes and their levels gradually increase as the oocyte approaches maturation near the spermatheca (Detwiler, Reuben, Li, Rogers, & Lin, 2001). As a result of the *oma-1/2* RNAi, the germline contained large defective oocytes that failed to complete maturation. Knockdown of *oma-1/2* resulted in increased GFP::MEX-3 expression in these oocytes (Fig. 2e, 2i, Table S4), confirming previous findings (Tsukamoto et al., 2017). Similarly, these results were also observed in the transgenic *mex-3* 3’UTR reporter strain (Fig. S1, Table S7). Therefore, OMA-1/2 appear to repress *mex-3* expression in the oocytes through the 3’UTR. Consistently, *mex-3* mRNA was found to associate with purified OMA-1 protein *in vitro* (Tsukamoto et al., 2017). The 3’UTR of *mex-3* contains clusters of OMA binding motifs UA(A/U). We previously demonstrated that OMA-1/2 bind to such motifs with a high degree of cooperativity (Kaymak & Ryder, 2013). Only *mex-3(spr9)* mutant animals showed increased GFP::MEX-3 expression in the oogenic region (Fig. 4b, 4c, Table S6). The 3’UTR region deleted in the *spr9* allele contains numerous UA(A/U) motifs (Fig. 4a). MEX-3 de-repression in the oocytes in the *mex-3(spr9)* mutant animals indicates that there are regions in the 3’UTR with high affinity to OMA-1/2.

Intriguingly, knockdown of *oma-1/2* caused a significant increase in the amount of GFP::MEX-3 in the oocyte nuclei. This suggests that OMA-1/2 plays a role, directly or indirectly, in MEX-3 partitioning between the nucleus and the cytoplasm in oogenesis. It remains unknown what role MEX-3 plays in the nucleus, if any, or if it actively shuttles between the two compartments.

Together, our candidate RBP RNAi results demonstrate that *mex-3* mRNA is post-transcriptionally regulated through its 3’UTR by different RNA-binding proteins throughout the germline (Fig. 6a). GLD-1 represses *mex-3* in the meiotic region. LIN-41 represses *mex-3* in the loop region and OMA-1/2 repress *mex-3* in the oocytes. Though not 3’UTR mediated, DAZ-1 represses *mex-3* in the distal mitotic end. Additionally, the largest 3’UTR deletion results in complete de-repression of MEX-3 expression throughout the germline including the mitotic end, meiotic region, loop region, and early oocytes. The expression patterns observed in the different mutants suggest that multiple RBP binding events contribute to complete repression in different regions of the germline.

### Polyadenylation and deadenylation play a role in regulating MEX-3 expression in the germline

Our results indicate that cytoplasmic polyadenylation contributes to the positive regulation of *mex-3* in the mitotic region and the oocytes. However, we do not yet know which RNA-binding partner is utilized by GLD-2 to bind and polyadenylate *mex-3* transcripts in the oocytes. We observed increased GFP::MEX-3 expression in all the stacked oocytes when we knocked down *gld-3* (Fig. 3c, 3h, Table S5). It is possible that all of these oocytes are all mature and that the stacking caused MEX-3 to condense. It is also possible that GFP::MEX-3 expression increased in the oocytes independent of the stacking and that this phenotype is an indirect result of *gld-3* knockdown. Since the binding specificity of GLD-3 is unknown, we do not know if the 3’UTR of *mex-3* contains any binding motifs for GLD-3. Our results show that polyadenylation contributes to the positive regulation of MEX-3 in the distal mitotic end and the oocytes (Fig. 6a). Our results also suggest a role for deadenylation in repressing *mex-3* in the meiotic region and mediating wild type expression of MEX-3 in the distal mitotic end. CCF-1 and NTL-1, but not CCR-4 appear to be essential components of this activity. These results support and expand upon a model where LIN-41 and OMA-1/2 repress their target mRNAs through the CCR4/NOT deadenylation complex during oocyte maturation (Tsukamoto et al., 2017).

### The translation initiation factor IFE-3 contributes to regulation of MEX-3 expression in the germline

Our results suggest that the translation initiation factor IFE-3 mediates *mex-3* repression in the germline. A previous study showed that IFE-3 confers a repressive effect mediated by its protein binding partners (Huggins et al., 2020). IFE-3 could form granules containing *mex-3* mRNA to inhibit its translation. In the meiotic region, IFE-3 may directly interact with *mex-3* transcripts to inhibit translation initiation. Alternatively, this interaction may be bridged by GLD-1, knockdown of which phenocopies loss of IFE-3 in the meiotic region. However, it has been shown by others that the translational efficiency of *gld-1* is reduced after *ife-3* knockdown ((Huggins et al., 2020), suggesting a third possibility that IFE-3 may indirectly repress *mex-3* by positively regulating a negative *mex-3* regulatory RBP. It will be interesting to further explore how exactly these mechanisms contribute to translational control of *mex-3* and other maternal mRNAs in the germline.

### An integrated model for coordination of MEX-3 expression and its contribution to fecundity

Our findings demonstrate that the 3’UTR of *mex-3* controls the unique spatiotemporal expression pattern of MEX-3 in the germline (Fig. 6a) and contributes to germline development and fecundity. Surprisingly, deleting the majority of the 3’UTR does not cause complete sterility but instead leads to reduced fecundity. This finding is reminiscent of several miRNA family mutations that are not essential under standard growth conditions, but are required during stressful conditions such as aging, exposure to pathogens, or growth at elevated temperatures (Brenner, Jasiewicz, Fahley, Kemp, & Abbott, 2010). For example, strong *let-7* loss of function mutant animals have a reduced lifespan when exposed to the pathogen *Pseudomonas aeruginosa* (Ren & Ambros, 2015). The 3’UTR of *mex-3* could act similarly to promote reproductive robustness under stressful growth conditions.

It remains unclear how overexpression of MEX-3 causes reduced fecundity. Increased MEX-3 concentration in the germline could lead to greater occupancy of sub-optimal MEX-3 binding motifs on both existing and new target transcripts. This may cause dysregulation of expression of those genes in the germline, reducing but not eliminating gamete production. It is also possible that fractions of MEX-3 could be sequestered into inactive granules, repressed through interactions with other proteins, or repressed through post-translational modifications. Consistent with this, some *mex-3(spr10)* mutant animals showed MEX-3 accumulation in granules in the defective oocytes (Fig. 5b). It will be interesting to investigate such post-translational regulatory mechanisms and how they play a role in controlling levels of MEX-3. And whether those mechanisms are redundant with 3’UTR-mediated regulation.

In the early embryo, MEX-3 localizes to both the anterior and posterior blastomeres. However, MEX-3 is only active in the anterior blastomere due to degradation of MEX-3 in the posterior blastomere. In the anterior, the RBPs MEX-5/6 are thought to bind and protect MEX-3. The degradation in the posterior blastomere is mediated by the RBP SPN-4 and the kinase protein PAR-4 (Huang & Hunter, 2015). SPN-4 is only expressed in late oocytes and the early embryo (Mootz et al., 2004; Ogura, Kishimoto, Mitani, Gengyo-Ando, & Kohara, 2003; Tsukamoto et al., 2017), but PAR-4 is expressed throughout the germline and in the early embryo (Watts, Morton, Bestman, & Kemphues, 2000). Therefore, PAR-4 could potentially mediate degradation of fractions of MEX-3 in the germline. Both post-transcriptional as well as post-translational regulatory mechanisms may contribute to MEX-3 expression pattern in the germline. However, we note that the 3’UTR is sufficient to pattern the expression of a reporter gene, so post-translational regulation through directed MEX-3 turnover may enforce the pattern of expression but is not absolutely required.

It will be intriguing to assess whether the endogenous 3’UTRs of other germline RBPs are equally dispensable for fertility. The majority of studies investigating germline RBPs function in *C. elegans* have relied on transgenic reporter strains (Elewa et al., 2015; Farley et al., 2008; Farley & Ryder, 2012; Hubstenberger, Cameron, Shtofman, Gutman, & Evans, 2012; Jeong, Verheyden, & Kimble, 2011; Merritt, Rasoloson, Ko, & Seydoux, 2008; Pagano et al., 2009). By targeting the endogenous 3’UTRs using CRISPR/Cas9, we can assess the importance of the UTR elements to biological function. This approach can also be applied in other organisms where key proteins exhibit unique spatial expression patterns to control early developmental processes.

## Materials and Methods

### Worm maintenance

All strains used were maintained by growing the animals on *E. coli* OP50 seeded NGM plates. N2 wild type strain was used as a control in all the brood size experiments. Each isolated mutant was outcrossed at least three times before analysis. Genotypes of all the strains in this paper are in table S1.

### RNAi

RNAi was performed by soaking animals in double-stranded RNA corresponding to the genomic cDNA sequence of the gene of interest. RNA was isolated from wild type N2 animals using trizol and phenol-chloroform extraction followed by RT-PCR using Superscript III One Step RT-PCR system with Platinum Taq DNA polymerase kit (ThermoFisher Scientific cat #: 12574026) to prepare the cDNA, which was used to amplify the template DNA used in the in vitro transcription (IVT) reaction to transcribe the dsRNA. Ambion MEGAscript T7 in vitro transcription kit (ThermoFisher Scientific cat #: AM1333) was used to prepare the dsRNA following the manufacturer’s protocol. The dsRNA was purified by phenol-chloroform extraction and isopropanol precipitation. The sequences of the oligos that were used to amplify the cDNA used as a template in the IVT reactions are in table S2. For the RNAi soaks, each tube contained 2µl of 5x soaking buffer, and 8µl of 500-1000ng/µl purified dsRNA. 0.5µl of M9 buffer containing arrested L1 animals was added to each individual tube. In the *lin-41* RNAi, L4 animals were placed in the dsRNA instead of L1s. The control tube contained 2µl of soaking buffer and 8µl of nuclease-free water. The soaked animals were incubated at 20°C or 25°C for 24 hours in the thermocycler. 20°C incubation temperature was used for the DG4269 ((tn1753[gfp::3xflag::mex-3]) strain while the 25°C temperature was used for the WRM24 (*sprSi17 [mex-5p::MODC PEST::GFP::H2B::mex-3 3’UTR + Cbr-unc-119(+)] II*) strain. After 24 hours, animals were placed on NGM plates seeded with *E. coli* OP50 and placed in the incubator. Once the animals reached adulthood, they were mounted on a 2% agarose pad on microscope slides, treated with 1mM levamisole to paralyze the animals, covered with a cover glass, then imaged.

### CRISPR/Cas9 mutagenesis

Ribonucleoprotein (RNP) mixes consisted of recombinant purified *Sp*Cas9 (final conc. = 2 µM), chemically synthesized crRNAs (final conc. = 40ng/µl) and tracrRNA (final conc. = 40ng/µl), commercial duplex buffer (30 mM HEPES, pH 7.5; 100 mM potassium acetate), and nuclease-free water. Sequences for the guide RNAs used are in table S3. *Sp*Cas9 was expressed from pET28a-Cas9-His (Addgene plasmid number 98158) and purified in our lab. The RNP mix was incubated at 37°C for 10 min. After the incubation, the plasmid pRF4 (*rol-6*) was added as a co-injection marker (final conc. = 50ng/µl). The mix was centrifuged at maximum speed for 5 min prior to loading a pulled borosilicate glass capillary injection needle. Young adult animals were microinjected in their gonads with the injection mix and then allowed to recover in M9 buffer on *E. coli* OP50 seeded NGM plates. The progeny of the injected animals was screened for the presence of roller animals, indicating a successful injection. All roller animals were singled out onto individual NGM plates, allowed to lay eggs, then lysed in a lysis buffer (30 mM Tris pH=8, 8 mM EDTA, 100 mM NaCl, 0.7% NP-40, 0.7% Tween-20 + proteinase K just prior to use). Lysates were frozen at −80°C for at least 10 min, then incubated at 65°C for 1 hour and 95°C for 15 min prior to genotyping PCR. For a 25µl PCR reaction, 2µl of the lysate was used as a template. The primers used to detect *mex-3* 3’UTR deletions were (forward primer: 5’-GGCGGAAACATGAATCTGAGCCC-3’, reverse primer: 5’-CGGACAATTGATCGGCCAATTGAC-3’). PCR reactions were run on a 1.5% TAE agarose gel. Single bands that are shorter than the wild type band indicate a homozygous mutation while two bands including the wild type band indicate a heterozygous mutation. Sanger sequencing of the purified PCR product was used to define the identity of the specific deletion.

### Poly(A) tail assay and TOPO cloning

N2, DG4269, and all *mex-3* mutant animals were collected and washed in M9 buffer then frozen in trizol and stored at −80°C. Total RNA was isolated from these animals using phenol-chloroform and isopropanol extraction. For the poly(A) tail assay, a poly(A) tail assay kit (ThermoFisher Scientific cat #: 764551KT) was used following the protocol outlined by the manufacturer. For the tail-specific primer set, a universal reverse primer provided in the kit was used for all the strains. For N2, DG4269, *mex-3(spr5), mex-3(spr6)*, and *mex-3(spr10)*, the forward primer 5’-CTACGCACAACTAACGGAGA-3’ was used. For *mex-3(spr9)*, the forward primer 5’-TCATGTCCTCCCTCAAAGG-3’ was used and for *mex-3(spr7)*, the forward primer 5’-CCCCAATATATATTCCTACAGTAGG-3’ was used. The PCR products were purified using a Zymo Research DNA clean and concentrator kit (cat #: D4034). The PCR products were cloned into a pCR™4-TOPO® TA vector using a TOPO TA Cloning kit (ThermoFisher Scientific cat #: K4575J10) following the manufacturer’s protocol. Plasmids containing the insert were analyzed using Sanger sequencing.

### Fluorescence microscopy

All of the imaging was done using a Zeiss Axioskop 2 plus microscope. ImageJ version 1.49 was used to quantify the images of the fluorescent animals from the RNAi experiments in the DG4269, WRM24, and the *mex-3* 3’UTR deletion mutant strains. For each animal, a line (width = 30 pixels) was drawn starting from the distal tip of the germline spanning the entire germline to the last oocyte. The fluorescence intensity was measured for each pixel in the line and then binned (total number = 20 bins for the DG4269 animals, 10 bins for the WRM24 animals). The fluorescence intensity from each animal was averaged across each bin. GraphPad Prism 7.04 was used to graph the mean fluorescence intensity for all the animals.

### Brood size

For each biological replicate, ∼25 individual L3/L4 animals were placed on individual NGM plates seeded with *E. coli* OP50. Each animal was moved to a fresh plate after two days initially, and then moved again daily until the completion of the experiment. The number of eggs and larvae on the plate, from which the animal was moved, was counted 1-2 days later. The number of progeny is the total number of eggs and larvae produced during the animal’s fertile period. All animals were grown and counted at 20°C. N2 wild type animals were used as the control. Each assay consisted of three biological replicates.

### Sterility assay

On day 1 for each strain, adult animals were bleached using 20% alkaline hypochlorite solution (final conc. 20% commercial bleach, 250 mM NaOH), washed 2x with M9 buffer and soaked in M9 buffer in a 1.7ml Eppendorf tube overnight. On day 2, the animals were placed on *E. coli* OP50 seeded NGM plates and incubated at 25°C. On day 3, animals were singled out onto individual plates and kept in the 25°C incubator. On day 5, each plate was scored for fertility or sterility based on the presence of viable progeny on the plate. Animals that did not have any viable progeny on the plate were scored as sterile.

### Data analysis

In the imaging studies, a two tailed student t-test was used to compare the mean fluorescence intensities. For RNAi conditions that were compared to the same control data, an unstacked one-way ANOVA was used to assess the overall significance. Post-hoc pairwise p-values were calculated using the Fisher’s LSD test then corrected for multiple hypotheses using a Bonferroni adjustment by multiplying the p-values by the number of hypotheses tested. To analyze the nuclear fluorescence intensity in the *oma-1/2* RNAi animals and controls, a circle with a radius of 15 pixels was drawn in the nucleus and another circle of the same radius drawn in the cytoplasm of the same oocyte. We calculated the ratio by dividing the nuclear fluorescence intensity by the cytoplasmic fluorescence intensity. We calculated the ratios for the two most proximal oocytes and then averaged the two ratios for each individual animal. A two-tailed student t-test was used to compare the ratios of the control and treated animals. Brood size data were analyzed using both Mann-Whitney U test and Kolmogorov-Smirnov nonparametric tests to compare the distributions between mutant and control strains. The data presented in each brood size assay represent a global analysis from three independent biological replicates. The p-values reported in figure 4 and supplemental figure 2 are from the Kolmogorov-Smirnov test.

## Acknowledgements

We thank all members of the Ryder lab for helpful discussions and reading of the manuscript. We also thank Dr. Victor Ambros for critical reading of the manuscript. This work is funded by an R01 NIH grant (5R01GM117237) to S.P.R.. The DG4269 strain was provided by the Caenorhabditis Genetics Center (CGC) which is funded by NIH Office of Research Infrastructure Programs (P40 OD010440).

## Author Contributions

M.A. performed all the experiments and analyses. M.A. and S.P.R. wrote and edited the manuscript.

