## Supplemental Figures and Tables for "Multiple RNA regulatory pathways coordinate the activity and expression pattern of a conserved germline RNA-binding protein"

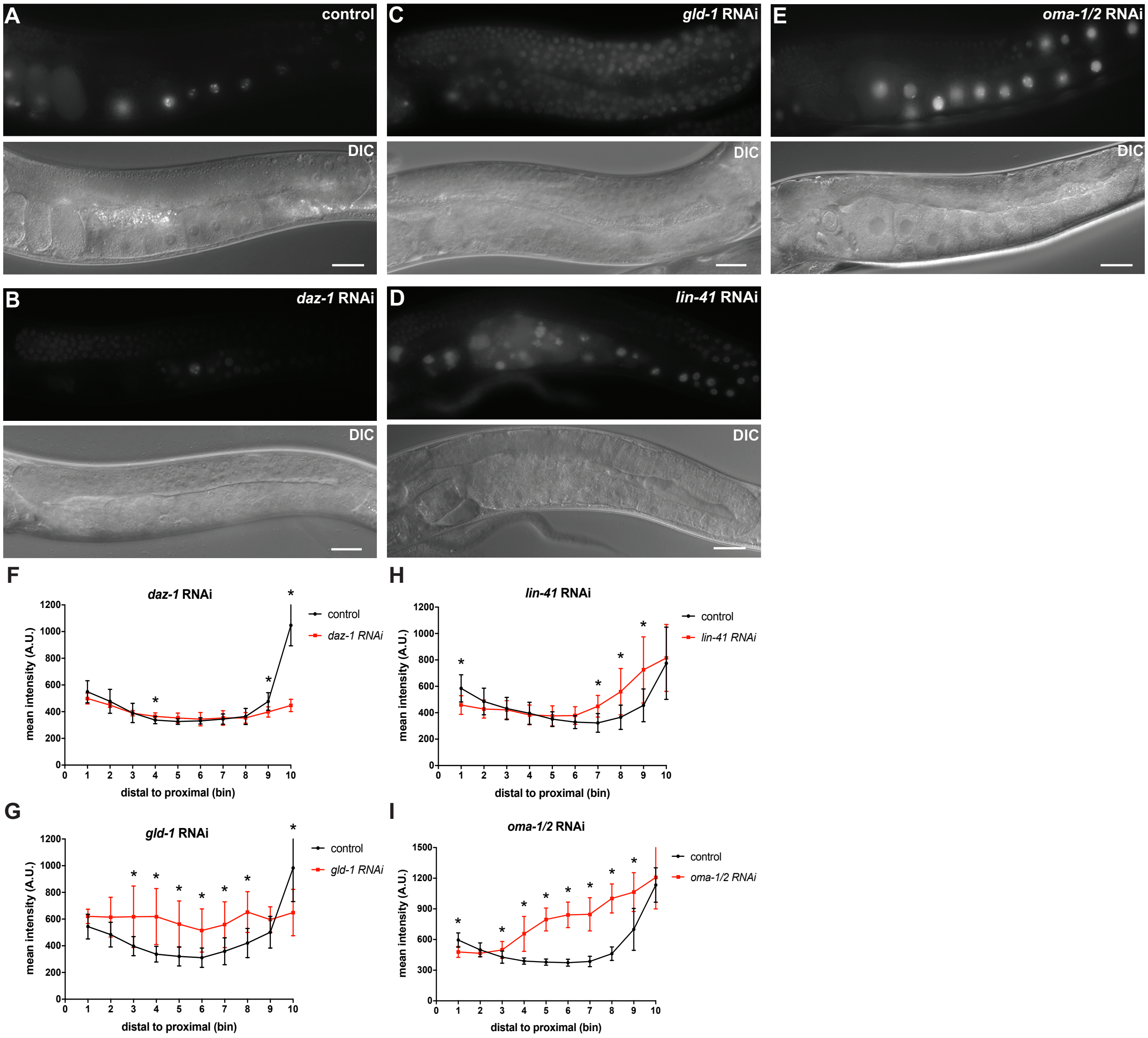


**Supplemental Figure 1. GLD-1, OMA-1/2, LIN-41, but not DAZ-1 regulate spatiotemporal expression pattern of MEX-3 through its 3´UTR. (A)** DIC and fluorescence images of *Pmex-5::MODC PEST::GFP::H2B::mex-3 3´UTR* animals from the control RNAi. **(B)** DIC and fluorescence images of *Pmex-5::MODC PEST::GFP::H2B::mex-3 3´UTR* animals after *daz-1* knockdown. *daz-1* knockdown didn´t cause increased reporter expression in the distal mitotic end. **(C)** DIC and fluorescence images of the transgenic reporter after *gld-1* knockdown. *gld-1* knockdown caused an overall increased expression of the reporter. **(D)** DIC and fluorescence of the transgenic reporter after *lin-41* knockdown. *lin-41* knockdown caused increased reporter expression in the loop region. **(E)** DIC and fluorescence images of transgenic reporter after *oma-1/2* knockdown. *oma-1/2* knockdown caused increased reporter expression in the oocytes. **(F)** quantitative analysis of the reporter fluorescence intensity after *daz-1* knockdown (n=10/10). **(G)** quantitative analysis of the reporter fluorescence intensity after *gld-1* knockdown (n=8/8). **(H)** quantitative analysis of the reporter fluorescence intensity after *lin-41* knockdown (n=8/8). **(I)** quantitative analysis of the reporter fluorescence intensity after *oma-1/2* knockdown (n=14/14). (*) indicates statistical significance, p-value ≤ 0.05. All p-values for this figure are reported in table S7. Scale bar = 30 µm.

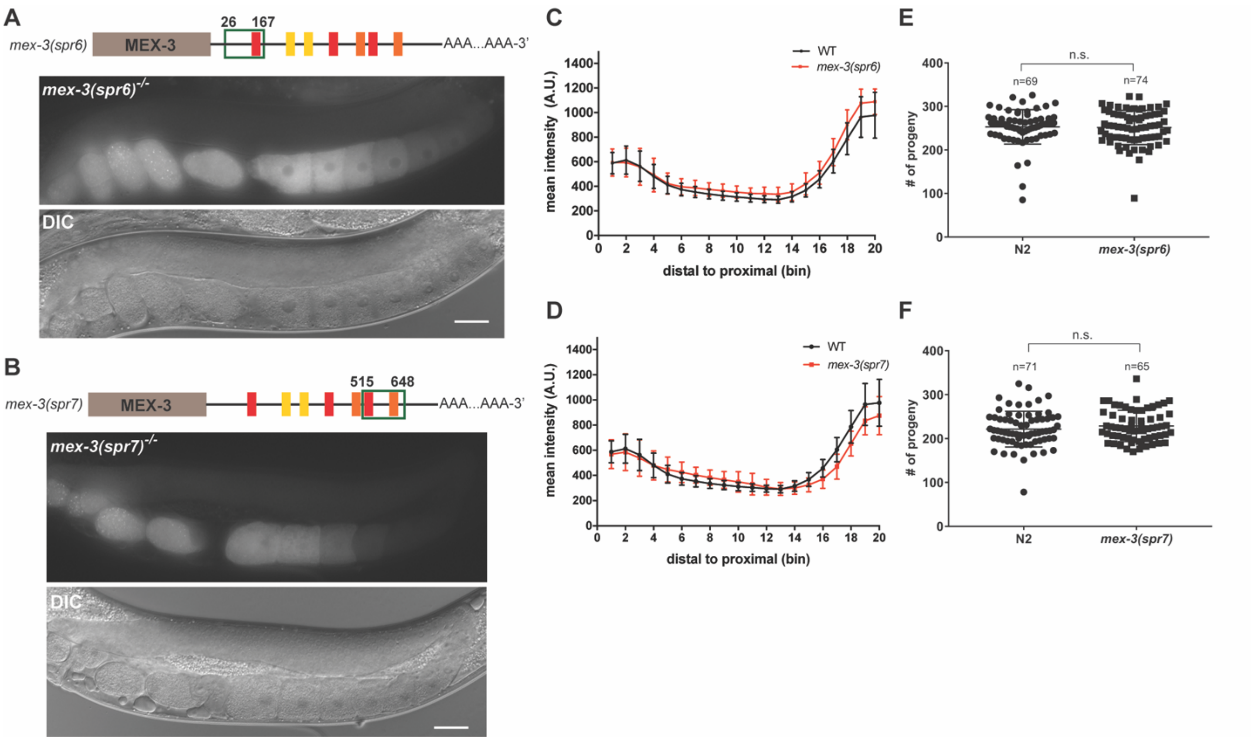


**Supplemental Figure 2. *mex-3* 3´UTR deletions in *mex-3(spr6)* and *mex-3(spr7)* did not alter MEX-3 expression or affect fertility**. **(A)** DIC and fluorescence images of the *mex-3(spr6)* homozygous mutant animals. **(B)** DIC and fluorescence images of the *mex-3(spr7)* homozygous mutant animals. **(C)** quantitative analysis of the fluorescence intensity in the *mex-3(spr6)* mutant animals compared to that of wild type GFP::MEX-3 (n=16). **(D)** quantitative analysis of fluorescence intensity in the *mex-3(spr7)* mutant animals compared to that of wild type GFP::MEX-3 (n=21). **(E)** brood size assay of *mex-3(spr6)* mutant animals at 20˚C.There is no reduction in the brood size. **(F)** brood size assay of *mex-3(spr7)* mutant animals at 20˚C. Each dot in panels E and F represents the brood size of an individual animal. In each panel, data from three biological replicates are shown in the graph. All p-values for panels C and D are reported in table S6. P-values in panels E and F are from a Kolmogorov-Smirnov test. All images taken at 40x magnification. Scale bar = 30 µm.

**Supplementary tables:**

**Table S1:** List of strains used in this paper

| Strain | Description |
|---|---|
| N2 | wild type |
| DG4269 | *mex-3(tn1753[gfp::3xflag::mex-3]) I* |
| WRM45 | *mex-3(spr5) I* |
| WRM49 | *mex-3(spr6) I* |
| WRM50 | *mex-3(spr7) I* |
| WRM52 | *mex-3(spr9) I* |
| WRM53 | *mex-3(spr10) I* |
| WRM24 | sprSi17 [mex-5p::MODC PEST::GFP::H2B::mex-3 3´UTR + Cbr-unc-119(+)] II |

**Table S2**: *In vitro* transcription primers for RNAi

| Name | Sequence |
|---|---|
| gld-1frag1F | 5´-ATGCCGTCGTGCACCACT-3´ |
| gld-1frag1R | 5´-CAATTGGCCTTTCCACGGTG-3´ |
| gld-2frag1F | 5´-GCAGAACGAAACGACGAACAC-3´ |
| gld-2frag1R | 5´-ATTGGATCCTGCCCGCGT-3´ |
| gld-3frag1F | 5´-ATGGGGGAGCAAAGCCATG-3´ |
| gld-3frag1R | 5´-TCCGGTGCCGGAGACTCG-3´ |
| oma-1frag1F | 5´-ATGAACGTTAACGGTGAAAACAACG-3´ |
| oma-1frag1R | 5´-GCGAGACGGTGGATAGGTCATC-3´ |
| oma-2frag1F | 5´-ATGGATATGCTCAAGGAAAATGTTATCC-3´ |
| oma-2frag1R | 5´-GTTGATGGATCCAACGGCCAG-3´ |
| lin-41frag1F | 5´- ATCGTGCCATGCTCATTGGAG-3´ |

| | |
|---|---|
| lin-41frag1R | 5´- GGAATTACGTGGTGGAGCTATGG-3´ |
| lin-41frag2F | 5´- GATGGCTACTTTGATGAGCCGT-3´ |
| lin-41frag2R | 5´- CTCACGGGGTGTCATTGTGAC-3´ |
| daz-1frag1F | 5´- ATGTCGCCGCCTCTACGGTATC-3´ |
| daz-1frag1R | 5´- CTGCTGCTGCTGCACATTGG-3´ |
| ccr-4fragF | 5´-GACAGTGGAGACGGCGAATC-3´ |
| ccr-4fragR | 5´-CCGGTGGAGGATGAGTGTTAACATG-3´ |
| ccf-1frag1F | 5´-ATGGCTTCTAGTAGCAGTGGTGG-3´ |
| ccf-1frag1R | 5´-GTCCACAGATCAATGGAGCAATCT-3´ |
| ntl-1fragF | 5´-CTCCAAAAAAGGCACCATCACGC-3´ |
| ntl-1fragR | 5´-CGGCGAATTATCCTGTCTTCCATTTTC-3´ |
| ntl-1frag2F | 5´-GGTGTGGAATGATACCAGCCTTTCAAAAC-3´ |
| ntl-1frag2R | 5´-GCCAGAGTTTGAATTCTGCCGTTGAGTAAG-3´ |
| ife-3fragF | 5´- GGGAGGATTGTCTGAAGATGGTTTCACTTTTCG-3´ |
| ife-3fragR | 5´- GAGATCTTTAGAATATGCTTAAGGAGTTGGGG-3´ |

**Table S3**: Guides RNAs used to target the *mex-3* 3´UTR using CRISPR/Cas9

| **Name** | **Sequence** |
|---|---|
| mex-3cr1 | 5´-GAGAGTCTACACGATAGTAA-3´ |
| mex-3cr2 | 5´-TATATATTGGGGGACTGCAT-3´ |
| mex-3cr3 | 5´-TAGTTGTGCGTAGTAGAGAG-3´ |
| mex-3cr4 | 5´-TTCACATACACCAAAATCTG-3´ |

**Table S4: P-values for bin to bin pairwise comparisons of mean fluorescence intensity in the GFP::MEX-3 strain in figure 2.** Adjusted p-values for *gld-1*, *daz-1*, and *lin-41* are corrected for multiple hypothesis testing as described in the methods, while p-values for *oma-1/2* are from a student t-test.

| **bin #** | ***gld-1*** | **Fold change** | ***daz-1*** | **Fold change** | ***lin-41*** | **Fold change** | ***oma-1/2*** | **Fold change** |
|---|---|---|---|---|---|---|---|---|
| 1 | 0.012 | 1.33 | 0.022 | 1.29 | 1.408 | 1.10 | 1.42E-02 | 1.22 |
| 2 | 0.004 | 1.37 | 0.012 | 1.31 | 4.304 | 1.03 | 0.4818 | 1.05 |
| 3 | 0.000 | 1.85 | 2.310 | 1.10 | 0.319 | 0.81 | 0.0598 | 0.85 |
| 4 | 0.000 | 2.50 | 3.838 | 0.95 | 0.151 | 0.78 | 0.0574 | 0.84 |
| 5 | 0.000 | 3.00 | 3.012 | 0.92 | 1.793 | 0.89 | 0.5329 | 0.94 |
| 6 | 0.000 | 3.37 | 4.385 | 0.96 | 3.805 | 1.05 | 0.1734 | 1.23 |
| 7 | 0.000 | 3.62 | 5.914 | 1.00 | 0.457 | 1.26 | 0.0217 | 2.05 |
| 8 | 0.000 | 3.52 | 5.046 | 1.05 | 0.054 | 1.54 | 4.50E-04 | 3.13 |
| 9 | 0.000 | 3.02 | 4.476 | 1.08 | 0.000 | 1.92 | 8.00E-05 | 3.53 |
| 10 | 0.000 | 2.57 | 4.081 | 1.12 | 0.000 | 2.21 | 4.86E-06 | 3.29 |
| 11 | 0.000 | 2.35 | 3.840 | 1.12 | 0.000 | 2.23 | 5.64E-07 | 2.92 |
| 12 | 0.001 | 2.29 | 4.657 | 1.08 | 0.000 | 2.05 | 7.15E-06 | 2.82 |
| 13 | 0.000 | 2.32 | 5.736 | 1.02 | 0.001 | 1.94 | 9.37E-07 | 2.91 |
| 14 | 0.000 | 2.47 | 4.611 | 0.92 | 0.001 | 1.88 | 4.37E-07 | 3.00 |
| 15 | 0.001 | 2.25 | 3.302 | 0.82 | 0.004 | 1.89 | 1.07E-08 | 2.96 |
| 16 | 0.000 | 1.88 | 0.607 | 0.68 | 0.000 | 1.79 | 8.06E-08 | 2.77 |
| 17 | 0.050 | 1.37 | 0.003 | 0.54 | 0.000 | 1.49 | 7.00E-05 | 1.93 |
| 18 | 4.490 | 1.04 | 0.000 | 0.42 | 0.628 | 1.16 | 0.0321 | 1.30 |
| 19 | 0.830 | 0.83 | 0.000 | 0.34 | 1.079 | 0.88 | 0.9039 | 0.99 |
| 20 | 0.016 | 0.67 | 0.000 | 0.32 | 0.003 | 0.70 | 0.0458 | 0.86 |

**Table S5: P-values for bin to bin pairwise comparisons of mean fluorescence intensity in the GFP::MEX-3 strain in figure 3.** Adjusted p-values for *ccf-1*, *ntl-1*, and *ife-3* are corrected for multiple hypothesis testing as described in the methods, while p-values for *gld-2* and *gld-3* are from a student t-test.

| bin # | *gld-2* | Fold change | *gld-3* | Fold change | *ccf-1* | Fold change | *ntl-1* | Fold change | *ife-3* | Fold change |
|---|---|---|---|---|---|---|---|---|---|---|
| 1 | 0.0273 | 0.81 | 0.8979 | 0.99 | 0.000 | 1.44 | 0.017 | 1.27 | 2.216 | 0.92 |
| 2 | 0.0078 | 0.82 | 0.9459 | 1.00 | 0.000 | 1.41 | 0.910 | 1.13 | 3.698 | 0.96 |
| 3 | 0.0003 | 0.77 | 0.5631 | 1.05 | 3.116 | 1.06 | 1.031 | 0.86 | 4.769 | 0.97 |
| 4 | 0.0019 | 0.81 | 0.3981 | 1.10 | 1.920 | 0.90 | 0.112 | 0.76 | 1.585 | 1.11 |
| 5 | 0.1225 | 0.90 | 0.3508 | 1.10 | 4.571 | 0.97 | 0.312 | 0.79 | 0.069 | 1.28 |
| 6 | 0.8322 | 0.99 | 0.0953 | 1.14 | 1.148 | 1.15 | 0.573 | 0.81 | 0.004 | 1.41 |
| 7 | 0.4870 | 1.05 | 0.0164 | 1.19 | 0.007 | 1.50 | 1.530 | 0.82 | 0.007 | 1.52 |
| 8 | 0.4606 | 1.05 | 0.0217 | 1.22 | 0.004 | 1.73 | 2.695 | 0.84 | 0.039 | 1.61 |
| 9 | 0.7759 | 1.02 | 0.0308 | 1.25 | 0.001 | 1.87 | 3.123 | 0.85 | 0.028 | 1.68 |
| 10 | 0.9706 | 1.00 | 0.0172 | 1.24 | 0.001 | 2.06 | 3.460 | 0.85 | 0.012 | 1.85 |
| 11 | 0.8608 | 0.99 | 0.0329 | 1.27 | 0.000 | 2.00 | 2.320 | 0.79 | 0.001 | 1.92 |
| 12 | 0.8790 | 0.99 | 0.0082 | 1.51 | 0.004 | 1.89 | 2.344 | 0.78 | 0.005 | 1.90 |
| 13 | 0.8904 | 0.99 | 0.0053 | 2.00 | 0.000 | 2.12 | 2.638 | 0.80 | 0.047 | 1.69 |
| 14 | 0.6629 | 0.97 | 0.0032 | 2.55 | 0.000 | 1.98 | 4.166 | 0.91 | 0.070 | 1.61 |
| 15 | 0.1421 | 0.91 | 0.0020 | 2.84 | 0.003 | 1.93 | 4.947 | 0.94 | 0.224 | 1.57 |
| 16 | 0.0339 | 0.84 | 0.0009 | 2.78 | 0.198 | 1.36 | 1.284 | 0.78 | 0.367 | 1.33 |
| 17 | 0.0406 | 0.81 | 0.0002 | 2.36 | 4.466 | 1.04 | 0.073 | 0.71 | 4.487 | 1.04 |
| 18 | 0.4110 | 0.92 | 0.0003 | 1.77 | 0.539 | 0.83 | 0.052 | 0.72 | 0.222 | 0.78 |
| 19 | 0.9340 | 1.01 | 0.0010 | 1.42 | 0.001 | 0.64 | 0.138 | 0.78 | 0.008 | 0.69 |
| 20 | 0.9242 | 1.01 | 0.0367 | 1.18 | 0.000 | 0.63 | 0.494 | 0.84 | 0.001 | 0.66 |

**Table S6: Student t-test p-values for bin to bin pairwise comparisons in the *mex-3* 3´UTR deletion mutants**

| bin # | *mex-3(spr6)* | Fold change | *mex-3(spr7)* | Fold change | *mex-3(spr9)* | Fold change | *mex-3(spr10)* | Fold change |
|---|---|---|---|---|---|---|---|---|
| 1 | 0.8821 | 1.01 | 0.6112 | 0.97 | 0.0163 | 1.18 | 0.238 | 0.93 |
| 2 | 0.6463 | 0.97 | 0.5451 | 0.95 | 0.0304 | 1.15 | 0.0374 | 0.87 |
| 3 | 0.9121 | 0.99 | 0.5995 | 0.96 | 0.0028 | 1.24 | 0.0633 | 0.88 |
| 4 | 0.7874 | 1.02 | 0.9839 | 1.00 | 4.91E-07 | 1.47 | 0.5389 | 0.96 |
| 5 | 0.6907 | 1.03 | 0.2443 | 1.09 | 3.87E-11 | 1.80 | 0.3872 | 1.05 |
| 6 | 0.3152 | 1.06 | 0.0452 | 1.14 | 0 | 2.17 | 0.159 | 1.07 |
| 7 | 0.1001 | 1.10 | 0.016 | 1.14 | 0 | 2.56 | 0.1092 | 1.08 |
| 8 | 0.0393 | 1.11 | 0.0159 | 1.14 | 0 | 2.87 | 0.065 | 1.10 |
| 9 | 0.0239 | 1.13 | 0.043 | 1.13 | 0 | 2.98 | 0.0671 | 1.11 |
| 10 | 0.0164 | 1.13 | 0.1226 | 1.12 | 0 | 3.01 | 0.0503 | 1.12 |
| 11 | 0.0102 | 1.13 | 0.2614 | 1.09 | 0 | 2.98 | 0.0494 | 1.13 |
| 12 | 0.0094 | 1.15 | 0.5356 | 1.04 | 0 | 2.80 | 0.0345 | 1.17 |
| 13 | 0.0157 | 1.15 | 0.8783 | 1.01 | 9.11E-11 | 2.48 | 0.017 | 1.24 |
| 14 | 0.0899 | 1.12 | 0.3192 | 0.95 | 2.28E-10 | 2.27 | 0.0218 | 1.28 |
| 15 | 0.0684 | 1.14 | 0.0328 | 0.88 | 1.26E-11 | 2.21 | 0.0265 | 1.35 |
| 16 | 0.0912 | 1.12 | 0.0015 | 0.81 | 7.52E-12 | 2.04 | 0.0383 | 1.32 |
| 17 | 0.0669 | 1.12 | 0.0004 | 0.78 | 7.52E-11 | 1.72 | 0.0817 | 1.20 |
| 18 | 0.0204 | 1.14 | 0.0016 | 0.83 | 2.11E-06 | 1.37 | 0.23 | 1.10 |
| 19 | 0.0382 | 1.12 | 0.0091 | 0.87 | 0.1166 | 1.10 | 0.8602 | 1.01 |
| 20 | 0.0498 | 1.11 | 0.0846 | 0.90 | 0.8395 | 0.99 | 0.6583 | 0.97 |

**Table S7: Student t-test p-values for bin to bin pairwise comparisons of mean fluorescence intensity in the *mex-3* 3´UTR transgenic reporter strain in supplemental figure 1**

| bin # | *gld-1* | Fold change | *daz-1* | Fold change | *lin-41* | Fold change | *oma-1/2* | Fold change |
|---|---|---|---|---|---|---|---|---|
| 1 | 0.1206 | 1.14 | 0.1026 | 0.91 | 0.0127 | 0.78 | 4.68E-05 | 0.80 |
| 2 | 0.0674 | 1.27 | 0.3354 | 0.94 | 0.2065 | 0.88 | 1.06E-01 | 0.93 |
| 3 | 0.0348 | 1.56 | 0.9630 | 1.00 | 0.8094 | 0.98 | 1.77E-02 | 1.17 |
| 4 | 0.0043 | 1.83 | 0.0415 | 1.08 | 0.7648 | 0.97 | 9.86E-06 | 1.68 |
| 5 | 0.0061 | 1.75 | 0.0803 | 1.08 | 0.4667 | 1.07 | 1.38E-12 | 2.10 |
| 6 | 0.0080 | 1.66 | 0.4952 | 1.04 | 0.1113 | 1.15 | 1.35E-12 | 2.25 |
| 7 | 0.0122 | 1.55 | 0.6957 | 1.03 | 0.0054 | 1.39 | 5.22E-10 | 2.20 |
| 8 | 0.0070 | 1.55 | 0.6804 | 0.97 | 0.0150 | 1.53 | 3.05E-12 | 2.17 |
| 9 | 0.0922 | 1.19 | 0.0048 | 0.83 | 0.0168 | 1.59 | 6.19E-05 | 1.52 |
| 10 | 0.0060 | 0.66 | 1.34E-09 | 0.43 | 0.7647 | 1.05 | 4.49E-01 | 1.07 |

**Table S8: The length of the *mex-3* 3´UTR from sequenced TOPO cloning plasmids containing tail-specific PCR products from the poly(A) tail assay. Each number represents a single plasmid sequenced**

| strain | 3´UTR Processing site |
|---|---|
| N2 | 684/687 |
| DG4269 | 685 |
| *mex-3(spr5)* | 685/684 |
| *mex-3(spr6)* | 685/684 |
| *mex-3(spr7)* | 685/685 |
| *mex-3(spr9)* | 685/685/687 |
| *mex-3(spr10)* | 687/641 |
